# Development of Dynamic Functional Architecture during Early Infancy

**DOI:** 10.1101/829846

**Authors:** Xuyun Wen, Rifeng Wang, Weili Lin, Han Zhang, Dinggang Shen

## Abstract

Understanding the moment-to-moment dynamics of functional connectivity (FC) in the human brain during early development is crucial for uncovering neuro-mechanisms of the emerging complex cognitive functions and behaviors. Instead of calculating FC in a static perspective, we leveraged a longitudinal resting-state functional magnetic resonances imaging dataset from fifty-one typically developing infants and, for the first time, thoroughly investigated how the temporal variability of the FC architecture develops at the *global* (entire brain), *meso*- (functional system) and *local* (brain region) levels in the first two years of age. Our results revealed that, in such a pivotal stage, *1)* the whole-brain FC dynamics is linearly increased; *2)* the high-order functional systems display increased FC dynamics for both within- and between-network connections, while the primary systems show the opposite trajectories; *3)* many frontal regions have increasing FC dynamics despite large heterogeneity in developmental trajectories and velocities. All these findings indicate that the brain is gradually reconfigured towards a more flexible, dynamic, and adaptive system with globally increasing but locally heterogeneous trajectories in the first two postnatal years, explaining why infants have emerging and rapidly developing high-order cognitive functions and complex behaviors.

## 1 Introduction

The first two postnatal years are a pivotal period of life with a rapid development in human brain structure (Knickmeyer et al., 2008; Tau and Peterson, 2010; Li et al., 2018) and function (Amsterdam, 1972; Haith et al., 1988; Reznick, 2008; Zhang et al., 2018; Wen et al., 2019) that could largely shape later behavioral and cognitive performance. Increasing evidence has been indicating that the basic human brain structural and functional framework is largely in shape by the end of two years old (Gilmore et al., 2018). Thus, understanding the developing patterns of the human brain during the first two postnatal years is critical and necessary. Yet, the related studies are still limited due to difficulties in image acquisition and analysis.

In recent years, resting-state functional magnetic resonance imaging (rs-fMRI) has been emerging as a useful tool for probing functional brain development (Smyser et al., 2010; Gao et al., 2011; Smyser et al., 2011; Gao et al., 2012a; Alcauter et al., 2014; Damaraju et al., 2014; De Asis-Cruz et al., 2015; Gao et al., 2015; Wen et al., 2019). The most popular approach for this type of study is to represent the infant brain as a complex network that consists of various brain regions as nodes linked by edges, estimated by functional connectivity (FC), and delineate developmental trajectories of the network topological properties across age (Smyser et al., 2010; Gao et al., 2011; Smyser et al., 2011; Gao et al., 2012a; Alcauter et al., 2014; Damaraju et al., 2014; De Asis-Cruz et al., 2015; Gao et al., 2015; Wen et al., 2019). Based on this method, existing studies have revealed the brain functional network development at multiple levels, including global (i.e., whole brain) (Gao et al., 2011; Wen et al., 2019), mesoscale (i.e., functional sub-networks) (Gao and Lin, 2012b; Gao et al., 2014; Gao et al., 2015;) and local levels (i.e., brain regions) (Wen et al., 2019). Generally, the brain network is globally reconfigured to be more efficiently balanced between local and global information communications from neonates to two years of age (Gao et al., 2011; Wen et al., 2019). Different functional sub-networks are developed with diverse developmental trajectories, with primary systems mature earlier than high-order function-related systems (Gao et al., 2014; Gao et al., 2015). From the local level, our recent findings suggest that more association regions gradually emerge as connector hubs facilitating information exchange and integration among functional sub-networks, while many primary regions become less centralized (Gao et al., 2011; Wen et al., 2019). Although providing valuable insights, these studies all investigate FC from a “*static*” point of view, ignoring its essential “*temporal dynamics*”.

Mountainous evidence has indicated that the brain functional network dynamically reconfigures itself over the course of seconds to minutes both at rest (Zalesky et al., 2014) and during task performance (Braun et al., 2015), supporting the adaptive integration and coordination between different neural systems in response to rapid changing internal and external stimulus (Calhoun et al., 2014; Hutchison et al., 2013). Hence, the investigation of the time-varying functional interactions provide insights into the understanding of how dynamics affects neural and behavioral adaptability (Zhang et al., 2016). More importantly, the analysis of temporal variation has more potential to capture sensitive changes in the healthy and diseased brain than the static FC analysis method (Rashid et al., 2016).

Characterizing FC dynamics during normative development could largely help to uncover neurodevelopmental mechanisms and understand behavioral changes. Emerging data demonstrate the time-varying properties of FC change across years of childhood to support a remarkable range of emerging cognitive abilities (Marusak et al., 2017). For instance, previous studies revealed age-related development in the frequency and dwelling time (how long the brain spends on a given mode) of specific FC “modes” (Allen et al., 2014; Hutchison and Morton, 2015; Qin et al., 2015; Marusak et al., 2017; Medaglia et al., 2018) and between-state transitions from late childhood to young adult (Allen et al., 2014; Qin et al., 2015). Such developmental changes in the dynamic FC are related to the reduced behavioral variability and more accurate performance (McIntosh et al., 2008). However, all existing evidence comes from the age groups spanning from late childhood (>7) to young adult (Allen et al., 2014; Hutchison and Morton, 2015; Qin et al., 2015; Marusak et al., 2017; Medaglia et al., 2018). As one of the most important periods of brain development, characterizing the time-varying FC in the first two years of life is highly desired. Unraveling the normative early development of whole-brain dynamic FC could also help early detection of developmental disorders, such as adolescence mental disorders (Knaus et al., 2008; Merikangas et al., 2010; Deco et al., 2015; Liu et al., 2015).

In this paper, leveraging longitudinal rs-fMRI from fifty-one typically developing infants, we used a recently developed novel dynamic FC approach to characterize the development of the temporal variation of FC profiles during the first two years of life (Zhang et al., 2016; Dong et al., 2018). Specifically, we evaluated the FC dynamics at multiple levels, ranging from the whole-brain, sub-network (Dong et al., 2018; Sun et al., 2018) to the region level (Zhang et al., 2016). We start from computing the variability of the whole-brain functional network, each functional sub-network (including within- and between-network variabilities) and each regional functional network, and then charted their respective developmental trajectories in the first two years of life. Based on previous findings, we hypothesize that, during the first two years of development, ***(1)*** the whole-brain variability is consistently increased due to the increasing flexibility of the brain functional connectome; ***(2)*** the high-order function-related sub-networks tend to display increased within-network FC temporal variability to support the increasing demand of flexible cognitive switching; ***(3)*** significantly increasing between-network variability more likely occurs between pair of high-order sub-networks for better adaptation and more sophisticated behaviors; and ***(4)*** brain regions with significantly increased variability are mainly located at the association areas, while those with decreased variability mainly distribute at the primary areas.

## 2 Materials and method

### 2.1 Subjects

Images were obtained from the subjects enrolled in the *Multi-visit Advanced Pediatric brain imaging study for characterizing structure and functional development (MAP Study)*. Study procedures were approved by the University of North Carolina at Chapel Hill Institutional Review Board and informed written consent were obtained from the parents of participants. After quality control, data of 51 normal developing infants with multiple longitudinal rs-fMRI scans, i.e., at 0 month (*N* = 29), 3 months (*N* = 26), 6 months (*N* = 32), 9 months (*N* = 31), 12 months (*N* = 31), 18 months (*N* = 31) and 24 months (*N* = 20), entered the analysis. The distribution of the ages of all included subjects was presented in **Fig. 1**. Detailed inclusion and exclusion criteria were described elsewhere (Gao et al., 2015).

**Fig. 1.**
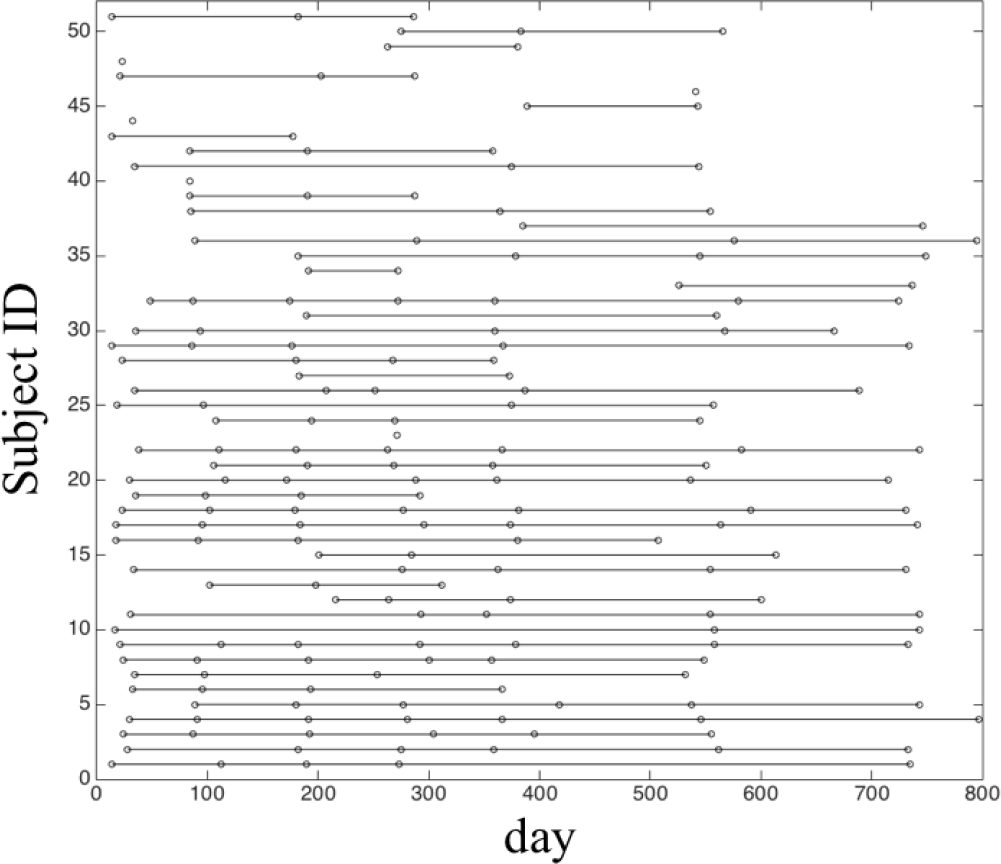
The distribution of the scans at different ages (days) for all the included subjects. Each dot represents one successful rs-fMRI scan from a subject at a certain age and the dots along each line represent all the available longitudinal scans of a subject.

### 2.2 Data Acquisition

During data acquisition, all subjects were in a natural sleeping state. No sedation was conducted. All subject data were acquired with a Siemens 3-T MR scanner. Rs-fMRI was acquired using T2-weighted EPI sequence with the following parameters: TR = 2 s, TE = 32 ms, 33 axial slices, voxel size of 4×4×4 mm^3^, and total volumes = 150 (5 min). Structural images were acquired with 3D MP-RAGE sequence with the following imaging parameters: TR = 1820 ms, TE = 4.38 ms, inversion time = 1100 ms, and voxel size = 1×1×1 mm^3^.

### 2.3 Data Preprocessing

Functional data were pre-processed using the tools from FMRIB Software Library (FSL, http://www.fmrib.ox.ac.uk/fsl). It includes the following steps: discarding the first 10 volumes, slice-timing correction, motion correction, and band-pass filtering (0.01 ~ 0.08 Hz). The mean signals from the white matter (WM), cerebrospinal fluid (CSF) and six motion parameters were removed using a linear regression model. To further reduce the head motion effects, wavelet despiking was used to remove both prolonged motion artifacts and higher frequency events (Patel et al., 2014). The subjects with the percentage of the detected spikes larger than 5% were excluded from further analysis.

Due to smaller and rapidly changing brain size and imaging contrast in the first two years of life, we used a specifically designed infant fMRI registration algorithm to align each data with a standard template in the Montreal Neuroscience Institute (MNI) space. To improve registration accuracy, we used tissue-label images, generated by a learning-based, infant-dedicated segmentation algorithm (Wang et al., 2015), for the following cross-sectional and longitudinal registration. First, we aligned each rs-fMRI data to its corresponding structural MRI data with linear registration. We then conducted a within-subject longitudinal registration that registers different structural images of the same subject at different ages to their “group mean” image using a group-wise registration method (Wu et al., 2012). After that, we adopted the Demons toolbox to align the group mean image of each subject to the standard symmetric “MNI-152” template (Thirion, 1998). By combining the linear transformation matrix and deformation fields from all above steps, we registered fMRI data of each subject at each scan from its native space to the common standard MNI space.

### 2.4 Temporal variability in FC Profile of Brain Regions and Whole Brain

To evaluate regional-wise variability of FC architecture, we first parcellated the whole brain into 268 regions by using an atlas in (Shen et al., 2013). As infant cerebellum registration is more difficult than the cerebral area registration due to its smaller size and weaker contrast, we excluded 51 cerebellar regions from further analysis. This resulted in 217 regions of interest (ROIs) in the cortical and subcortical areas. For each ROI, we extracted its mean time series by averaging the signals of all voxels within it and segmented the time course into *N* non-overlapping windows with an equal length of *L*. In the *i*-th window, we computed pairwise Pearson’s correlation among each pair of ROIs (i.e., nodes) using the windowed time series, generating a *M* × *M* FC matrix (denoted by *F*_*i*_, *i* = 1, 2, 3, …, *n*; *M* is the total number of ROIs; here, *M* = 217). The *k*-th row (or *k-*th column) in *F*_*i*_ characterizes the *FC* architecture for the ROI *k* at the *i*-th time window, represented as 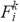. Then, the regional temporal variability of the ROI *k* (i.e., 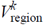) was defined as,

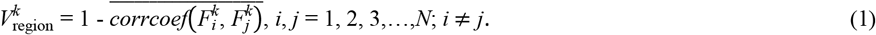

In Eq. 1, 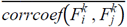 measures the averaged similarity of the FC architecture of the region *k* between any two different time windows and *corrcoef* denotes Pearson’s correlation coefficient. By deducting from 1, 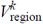 indicates the temporal variability of FC architecture of the region *k* (Zhang et al., 2016). To avoid the arbitrary choice of the window length and starting point of the window, *V*_region_ was computed with different window lengths (*L* = 10, 11,…, 20 time points) using *L*-1 different starting points and then averaged cross all window length and starting point parameters as the final regional dynamic FC variability.

After obtaining 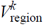, we calculated the whole-brain temporal variability of FC, *V*_whole-brain_, for each scan of each subject by averaging the 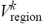 across all the 217 ROIs.

### 2.5 Variability of FC Architecture within and between Functional Sub-networks

To measure the mesoscale temporal variability of the FC architecture, we divided 217 ROIs into seven predefined functional sub-networks, each of which represents a certain functional system (Yeo et al., 2014). They include visual network (VN), sensorimotor network (SMN), dorsal attention network (DAN), ventral attention network (VAN), frontal-parietal network (FPN), default mode network (DMN), and limbic/striatal network (LN/SN) (Yeo et al., 2014). Because there are two major types of FC in terms of sub-network affiliations of two nodes that form a link, for each functional sub-network, we computed the within-network temporal variability of FC (*V*_within-net_), and between-network temporal variability of FC (*V*_between-net_) according to a modified algorithm of the regional temporal variability method of FC architecture (Dong et al., 2018). Specifically, 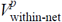 is defined as the time-varying dynamic of the FC architecture within functional sub-network *p* according to Eq. 2, and 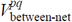 as the time-varying dynamics of the FC architecture between functional sub-networks *p* and *q* according to Eq. 3.

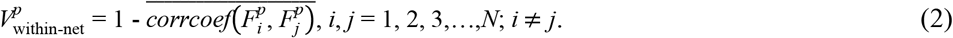

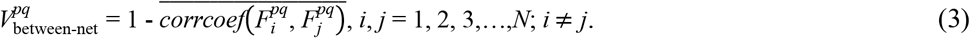

In Eqs. 2 and 3, *p* and *q* are indexes of seven functional sub-networks. 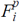 is the FC architecture consisting of all FCs in functional sub-network *p* at the *i*-th time window. 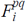 represents the FC architecture including all FCs between sub-networks *p* and *q* at the *i*-th time window. Likewise, we computed *V*_within-net_ and *V*_between-net_ with multiple window lengths (*L* = 10, 11, …, 20 time points) using different starting points and took an average to generate the final version of the *V*_within-net_ and *V*_between-net_.

### 2.5 Characterization of Developmental Trajectories of Variability

The developmental trajectories of the temporal variability of FC architecture at the whole-brain, sub-network, and regional levels were delineated by using linear mixed effect regression (LMER). The LMER was used due to its ability to handle missing data in longitudinal studies (Verbeke, 1997). For each measurement (*V*_whole-brain_, *V*_within-net_, *V*_between-net_, and *V*_region_), both a linear model (with age as a fixed effect variable) and a log-linear model (with log(age) as a fixed effect variable) were built. The log-linear model was used to delineate the uneven developmental speed during the first two years of life, which has been identified to follow the exponential function (Gao W et al., 2015). In each LMER model, *V*_whole-brain_, *V*_within-net_, *V*_between-net_, or *V*_region_ is the dependent variable, and age or log(age) in days is the independent variable. Subject-specific intercept and slope were included as random effect variables. Akaike information criterion (AIC) was used to gauge the selection between the linear and log-linear models. Significant developmental changes were determined by *p* < 0.05 after multiple comparison correction by false discovery rate (FDR).

## 3 Results

### 3.1 Temporal Variability of Whole-brain FC Architecture

The temporal variability of the whole-brain FC architecture was found significantly increased in a linear manner during the first two years of life (*p* < 0.05), suggesting that the brain becomes more and more flexible along development (Fig. 2a).

**Fig. 2.**
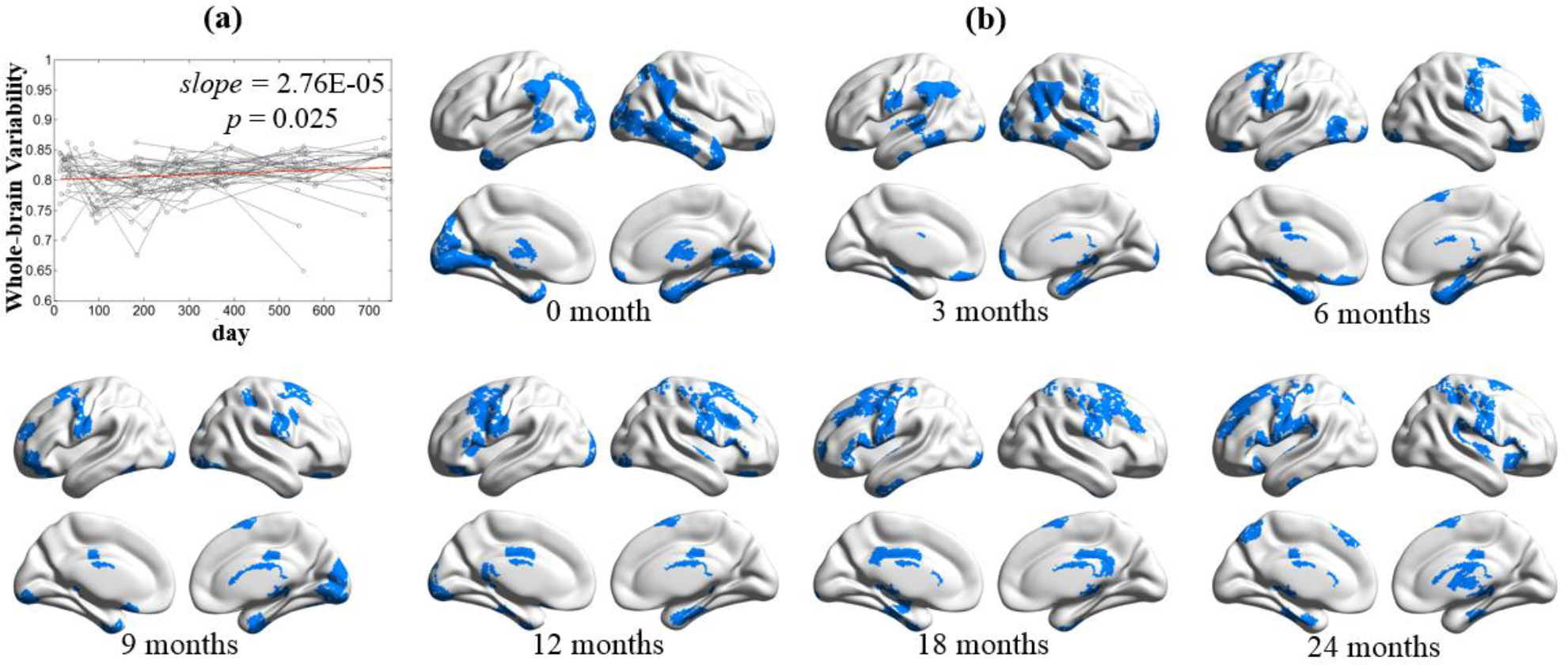
(a) Developmental trajectory of the temporal variability of whole-brain FC architecture across age. (b) Spatial patterns of 20% brain regions with the highest variability across seven age groups (i.e., 0, 3, 6, 9, 12, 18 and 24 months of age).

To clearly show spatial pattern changes at different ages, especially the regions with high FC variability, we further visualized the top 20% of the regions in terms of age-averaged regional FC temporal variability for the ages of 0, 3, 6, 9, 12, 18, 24 months (Fig. 2b). In general, we found that the spatial pattern of the high FC temporal variability regions is shifted from the posterior to the anterior of the brain, especially from the occipital and temporal lobes to the frontal lobe. Specifically, for neonates, brain regions with highly variable FC profile are mainly located at visual cortex, middle and inferior temporal cortices, and thalamus. After 3 months of development, those in the visual cortex, temporal cortex, and thalamus shrink largely, while bilateral sensorimotor areas and angular areas emerge. From 6 to 9 months old, the sensorimotor areas have continuously increased FC variability, while those in other regions are gradually weakened. Since 12 months of age, the spatial distribution of the top variable brain regions tends to be stable with a continuous pattern in bilateral sensorimotor areas but with prominent extension to the premotor area and superior frontal areas.

### 3.2 Temporal Variability of Sub-network’s FC Architecture

Fig. 3 plotted the developmental patterns of within-network and between-network FC temporal variability for seven functional sub-networks, with Table 1 summarizing the fitted developmental trajectories from the LMER model. One striking result is that two primary sub-networks (i.e., VN and SMN) showed exponentially decreased *V*_within-net_, while the *V*_within-net_ on other two high-order function-related sub-networks (i.e., VAN and DMN) are linearly increased. Such a finding mirrors different developmental reconfigurations on FC dynamics between primary sensory and high-level functional systems.

**Fig. 3.**
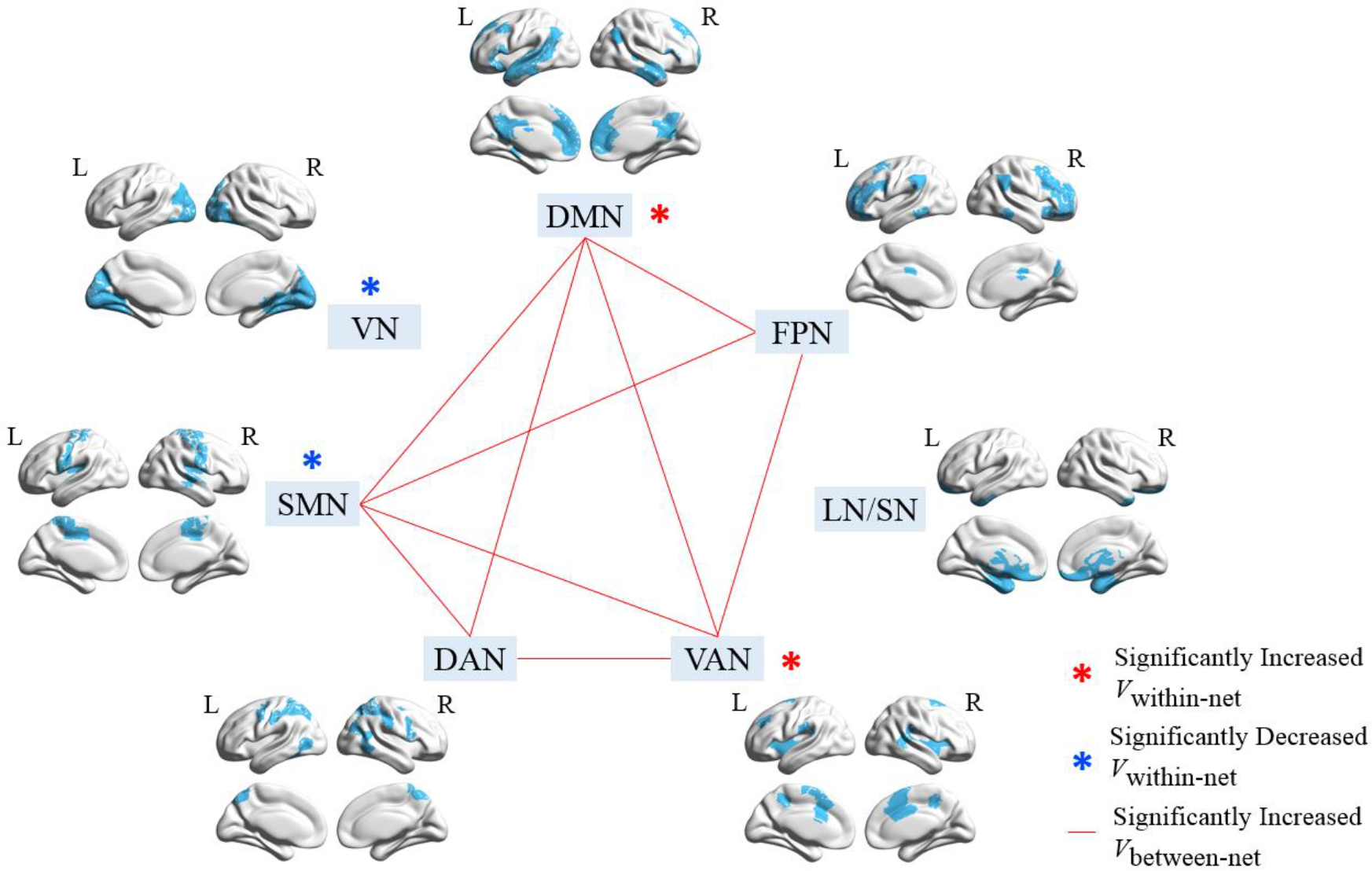
Illustration of developmental patterns of within-network (*V*_within-net_) and between-network FC temporal variabilities (*V*_between-net_) on seven functional sub-networks, including visual network (VN), sensorimotor network (SMN), dorsal attention network (DAN), ventral attention network (VAN), limbic/striatum network (LN/SN), default mode network (DMN), and frontal-parietal network (FPN). The sub-network parcellation is based on the template provided by Yeo et al (2011). Blue and red stars indicate significantly decreased and increased *V*_within-net_, respectively. The red lines between two sub-networks indicate significantly increased *V*_between-net_ between them. The significance level is *p* < 0.05 after FDR correction.

**Table 1.** Fitted Developmental Trajectories of Within-network Variability and Between-network Variability

| Network | Model | Direction | Intercept | Slope | *p* (Slope) | AIC Linear | AIC Log-linear |
| --- | --- | --- | --- | --- | --- | --- | --- |
| **Within-network Variability** |  |  |  |  |  |  |  |
| **VN** | Log-linear | ↓ | 0.87762 | -0.02731 | 4.76E-08 | -512.76 | -517.82 |
| **SMN** | Log-linear | ↓ | 0.81340 | -0.01777 | 3.48E-05 | -534.40 | -549.25 |
| DAN | Log-linear |  | 0.78918 | -0.00444 | 2.89E-01 | -562.33 | -565.97 |
| **VAN** | Linear | ↑ | 0.71142 | 0.00006 | 9.30E-03 | -471.89 | -465.78 |
| LN/SN | Linear |  | 0.77696 | 0.00001 | 3.88E-01 | -615.67 | -615.12 |
| FPN | Linear |  | 0.76788 | 0.00002 | 4.05E-01 | -498.01 | -494.54 |
| **DMN** | Linear | ↑ | 0.75920 | 0.00006 | 1.08E-03 | -610.51 | -599.80 |
| **Between-network Variability** |  |  |  |  |  |  |  |
| VN – SMN | Log-linear |  | 0.91685 | -0.00199 | 5.87E-01 | -595.46 | -595.85 |
| VN – DAN | Log-linear |  | 0.87219 | -0.00677 | 6.50E-02 | -617.93 | -619.10 |
| VN – VAN | Linear |  | 0.88072 | 0.00004 | 2.82E-02 | -580.80 | -578.07 |
| VN – LN/SN | Log-linear |  | 0.85471 | 0.00432 | 2.29E-01 | -673.85 | -677.25 |
| VN – FPN | Linear |  | 0.88661 | -0.00001 | 6.82E-01 | -612.80 | -600.87 |
| VN – DMN | Linear |  | 0.85318 | 0.00003 | 6.04E-02 | -629.80 | -627.42 |
| **SMN – DAN** | Linear | ↑ | 0.78940 | 0.00005 | 1.14E-02 | -537.27 | -536.65 |
| **SMN – VAN** | Linear | ↑ | 0.76950 | 0.00007 | 5.11E-04 | -584.01 | -581.32 |
| SMN – LN/SN | Linear |  | 0.89790 | 0.00002 | 1.62E-01 | -642.25 | -642.10 |
| **SMN – FPN** | Log-linear | ↑ | 0.82378 | 0.01049 | 1.80E-03 | -609.98 | -612.15 |
| **SMN – DMN** | Linear | ↑ | 0.84190 | 0.00006 | 3.43E-04 | -615.21 | -609.31 |
| **DAN – VAN** | Linear | ↑ | 0.81193 | 0.00006 | 9.53E-04 | -564.33 | -558.59 |
| DAN – LN/SN | Log-linear |  | 0.90896 | -0.00474 | 2.48E-01 | -645.76 | -653.29 |
| DAN – FPN | Log-linear |  | 0.78006 | 0.00899 | 1.81E-02 | -597.21 | -598.20 |

Note: VN: visual network; SMN: sensorimotor network; DAN: dorsal attention network; VAN: ventral attention network; LN/SN: subcortical network/limbic network; FPN: frontal-parietal network; DMN: default mode network. Statistically significant increases and decreases of FC temporal variability with age are shown with ↑ and ↓, respectively, and the corresponding network labels are shown in bold.

**Table 1 (Cont'd)** Fitted Developmental Trajectories of Within-network Variability and Between-network Variability
| Network | Model | Direction | Intercept | Slope | *p* (Slope) | AIC Linear | AIC Log-linear |
| --- | --- | --- | --- | --- | --- | --- | --- |
| **Between-network Variability** |  |  |  |  |  |  |  |
| **DAN – DMN** | Linear | ↑ | 0.82032 | 0.00007 | 4.65E-05 | -606.00 | -593.86 |
| VAN – LN/SN | Linear |  | 0.83864 | 0.00005 | 6.35E-02 | -553.30 | -547.53 |
| **VAN – FPN** | Linear | ↑ | 0.80804 | 0.00007 | 8.52E-05 | -629.37 | -626.97 |
| **VAN – DMN** | Linear | ↑ | 0.78999 | 0.00006 | 4.80E-03 | -583.16 | -573.53 |
| LN/SN – FPN | Linear |  | 0.85837 | 0.00002 | 3.56E-01 | -641.75 | -637.79 |
| LN/SN – DMN | Linear |  | 0.83016 | 0.00002 | 1.94E-01 | -635.65 | -633.73 |
| **FPN – DMN** | Linear | ↑ | 0.79962 | 0.00006 | 9.58E-05 | -633.02 | -630.32 |

Note: VN: visual network; SMN: sensorimotor network; DAN: dorsal attention network; VAN: ventral attention network; LN/SN: subcortical network/limbic network; FPN: frontal-parietal network; DMN: default mode network. Statistically significant increases and decreases of FC temporal variability with age are shown with ↑ and ↓, respectively, and the corresponding network labels are shown in bold.

For between-network FC temporal variability, we found all significantly developmental changes (FDR correction, *p*<0.05) are increased with age (Fig. 3). Eight of them showed the linear increase and one showed the exponential (i.e., nonlinear) increase. Interestingly, there is no *V*_between-net_ increases between VN (and LN/SN) and any other functional sub-networks. For other five functional sub-networks, all their *V*_between-net_ are found to increase with age. Specifically, for the SMN and DMN, more than half of their connections (4 out of 6) have the increased *V*_between-net_; and for DAN and FPN, half of their connections (3 out of 6) have the increased *V*_between-net_. Notably, except SMN, all the other networks with significantly increased *V*_between-net_ are the high-level functional networks (DMN, DAN, VAN, and FPN).

### 3.3 Temporal Variability of Regional FC Architecture

In addition to the above results at the global (i.e., whole brain) and sub-network levels, we also delineated developmental trajectories of variabilities for all the brain regions (ROIs). Of all 217 ROIs, 70 showed significant changes along the development (*p* < 0.05 after FDR correction), as visualized in Fig. 4a. Most (58/70, 82%) of the changes are increases. For the 58 ROIs with increasing trajectories, most of them are linear increases. We further classified these brain regions into four categories based on the types of their developmental trajectories, i.e., linear increase (Type 1), nonlinear increase (Type 2), linear decrease (Type 3), and nonlinear decrease (Type 4). See Fig. 4 and Table 2 for details.

**Fig. 4.**
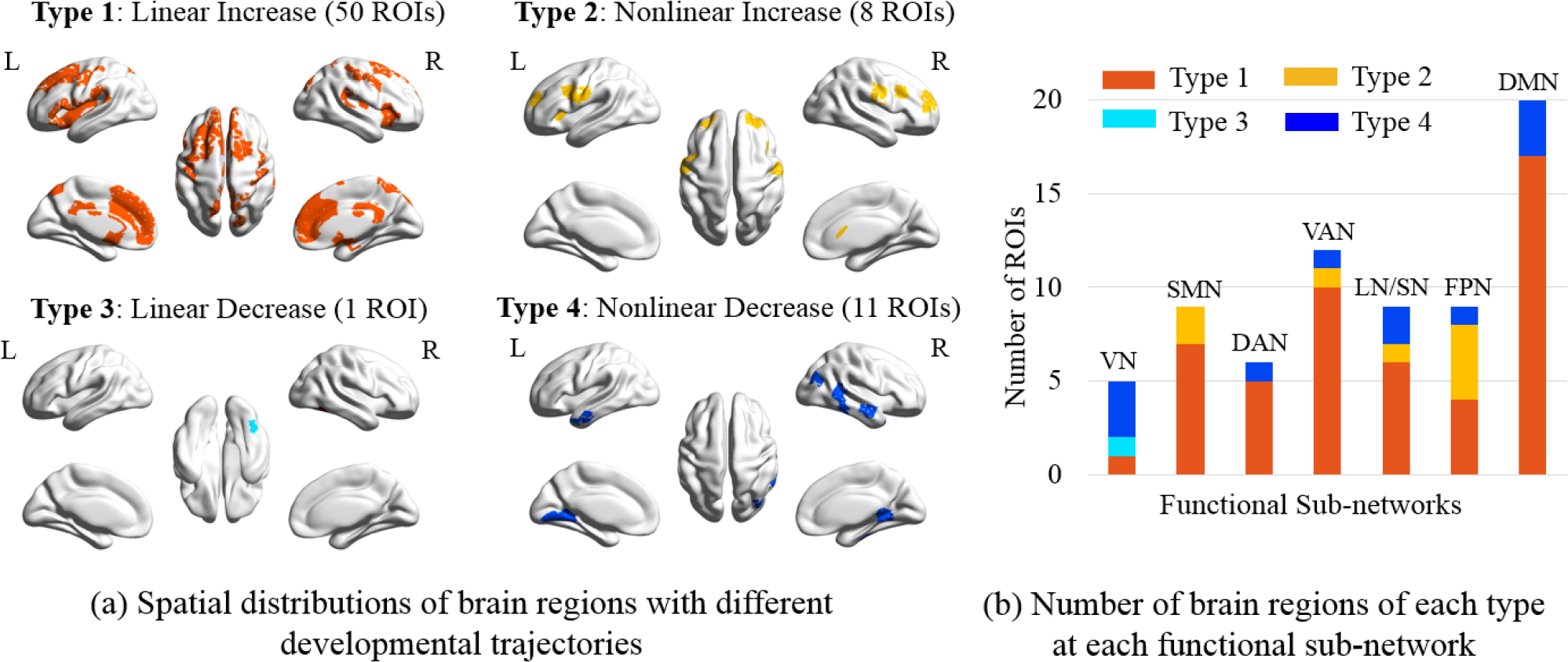
Four different categories of brain regions with significant development of FC temporal variability in the first two years of life. Type 1 and Type 2 respectively represent the brain regions with linear and nonlinear increases in variability. Type 3 and Type 4 are the regions with linear and nonlinear decreases in variability. (a) Spatial distributions of brain regions of four categories (denoted in different colors). (b) The number of brain regions of each category located in seven functional brain sub-networks, including visual network (VN), sensorimotor network (SMN), dorsal attention network (DAN), ventral attention network (VAN), limbic/striatal network (LN/SN), frontal-parietal network (FPN), and default mode network (DMN).

**Table 2.** Fitted Developmental Trajectories of Regional Variability

| Brain Region | Intercept | Slope | *p* (Slope) | AIC (Linear) | AIC (Log-linear) |
| --- | --- | --- | --- | --- | --- |
| Type 1: Linear Increase |  |  |  |  |  |
| Frontal_Med_Orb_R | 0.75028 | 0.00009 | 3.19E-04 | -491.79 | -483.48 |
| Frontal_Sup_Medial_R | 0.75566 | 0.00008 | 1.77E-03 | -440.78 | -437.45 |
| Frontal_Sup_R | 0.76529 | 0.00009 | 6.83E-04 | -460.24 | -453.29 |
| Cingulum_Mid_R | 0.73748 | 0.00011 | 2.66E-05 | -453.47 | -438.45 |
| Frontal_Inf_Oper_R | 0.77559 | 0.00010 | 9.17E-04 | -416.22 | -412.72 |
| Frontal_Inf_Tri_R | 0.75504 | 0.00012 | 8.55E-05 | -407.13 | -401.58 |
| Precentral_R | 0.81716 | 0.00006 | 4.64E-03 | -482.05 | -479.04 |
| Frontal_Sup_R | 0.81301 | 0.00008 | 7.80E-04 | -519.56 | -519.42 |
| Frontal_Sup_R | 0.80073 | 0.00007 | 6.27E-03 | -441.01 | -439.09 |
| Precentral_R | 0.80037 | 0.00007 | 8.79E-03 | -432.61 | -430.26 |
| Precentral_R | 0.79557 | 0.00010 | 1.30E-05 | -494.73 | -488.37 |
| Frontal_Inf_Orb_R | 0.73955 | 0.00015 | 1.69E-06 | -361.39 | -350.17 |
| Rolandic_Oper_R | 0.77817 | 0.00008 | 2.63E-03 | -450.67 | -450.65 |
| Parietal_Sup_R | 0.79223 | 0.00009 | 3.63E-04 | -483.48 | -471.59 |
| Precuneus_R | 0.77816 | 0.00008 | 4.65E-04 | -459.37 | -450.68 |
| Temporal_Sup_R | 0.78125 | 0.00010 | 1.70E-05 | -475.89 | -464.94 |
| Cuneus_R | 0.78537 | 0.00007 | 1.78E-03 | -476.74 | -470.38 |
| Cingulum_Mid_L | 0.71981 | 0.00016 | 2.04E-09 | -487.68 | -461.02 |
| Cingulum_Post_R | 0.79399 | 0.00005 | 1.26E-02 | -492.81 | -489.14 |
| Cingulum_Mid_R | 0.80329 | 0.00010 | 1.96E-05 | -489.87 | -483.59 |
| Hippocampus_R | 0.77957 | 0.00007 | 1.14E-02 | -428.05 | -427.08 |
| Cingulate Gyrus | 0.83407 | 0.00008 | 2.76E-04 | -494.10 | -492.61 |
| Putamen_R | 0.76433 | 0.00008 | 6.39E-03 | -423.95 | -421.11 |
| Frontal_Med_Orb_L | 0.77231 | 0.00007 | 2.37E-03 | -489.09 | -483.02 |
| Frontal_Sup_Medial_L | 0.73909 | 0.00012 | 9.39E-07 | -484.31 | -468.50 |
| Frontal_Sup_Medial_L | 0.77546 | 0.00009 | 1.88E-04 | -459.26 | -457.49 |
| Frontal_Mid_L | 0.77683 | 0.00012 | 3.74E-06 | -456.45 | -452.85 |

Note: We named each ROI using a traditional naming convention. The same brain ROI names in the table locate at different places, representing different sub-divisions of the corresponding (larger) brain regions.

**Table 2 (Cont'd)** Fitted Developmental Trajectories of Regional Variability
| Brain Region | Intercept | Slope | *p* (Slope) | AIC (Linear) | AIC (Log-linear) |
| --- | --- | --- | --- | --- | --- |
| Type 1: Linear Increase |  |  |  |  |  |
| Frontal_Sup_Medial_L | 0.76533 | 0.00011 | 1.03E-05 | -450.78 | -437.29 |
| Frontal_Mid_L | 0.78228 | 0.00010 | 9.01E-06 | -479.01 | -475.61 |
| Supp_Motor_Area_L | 0.74923 | 0.00010 | 7.91E-04 | -423.87 | -411.32 |
| Frontal_Inf_Orb_L | 0.76851 | 0.00013 | 4.68E-06 | -414.37 | -414.12 |
| Frontal_Inf_Orb_L | 0.74317 | 0.00016 | 3.89E-07 | -381.91 | -368.34 |
| Frontal_Inf_Tri_L | 0.76153 | 0.00015 | 2.93E-08 | -428.55 | -423.20 |
| Postcentral_L | 0.78182 | 0.00009 | 1.25E-03 | -457.69 | -453.03 |
| Temporal_Sup_L | 0.78590 | 0.00009 | 1.51E-04 | -455.11 | -452.18 |
| Frontal_Sup_L | 0.79952 | 0.00009 | 4.24E-04 | -429.98 | -424.26 |
| Precentral_L | 0.80872 | 0.00010 | 2.22E-05 | -506.05 | -504.44 |
| Postcentral_L | 0.77635 | 0.00012 | 2.74E-05 | -445.05 | -437.70 |
| Rolandic_Oper_L | 0.75287 | 0.00009 | 4.45E-03 | -425.29 | -418.22 |
| Insula_L | 0.74483 | 0.00010 | 7.35E-04 | -384.33 | -377.21 |
| Rolandic_Oper_L | 0.76468 | 0.00009 | 3.86E-04 | -437.53 | -434.76 |
| Parietal_Sup_L | 0.77447 | 0.00010 | 1.07E-04 | -450.91 | -438.99 |
| Temporal_Sup_L | 0.78964 | 0.00008 | 1.60E-03 | -460.51 | -455.99 |
| Cingulate Gyrus | 0.82975 | 0.00007 | 2.98E-03 | -472.26 | -471.92 |
| Cingulum_Ant_L | 0.71996 | 0.00015 | 3.89E-07 | -449.69 | -431.68 |
| Cingulum_Ant_L | 0.75333 | 0.00008 | 4.72E-04 | -459.96 | -451.82 |
| Cingulum_Mid_L | 0.78559 | 0.00007 | 9.84E-04 | -506.77 | -499.21 |
| Cingulum_Mid_L | 0.80448 | 0.00009 | 1.76E-04 | -485.13 | -480.85 |
| Cingulum_Ant_L | 0.76947 | 0.00013 | 1.14E-06 | -506.85 | -491.81 |
| Caudate_L | 0.75855 | 0.00008 | 4.65E-03 | -433.74 | -430.66 |
| Type 2: Nonlinear Increase |  |  |  |  |  |
| Frontal_Mid_R | 0.71852 | 0.01729 | 1.40E-03 | -454.39 | -459.20 |
| Frontal_Inf_Tri_R | 0.70181 | 0.01939 | 3.45E-04 | -464.38 | -465.10 |
| Postcentral_R | 0.73503 | 0.01937 | 1.06E-04 | -491.16 | -493.24 |

Note: We named each ROI using a traditional naming convention. The same brain ROI names in the table locate at different places, representing different sub-divisions of the corresponding (larger) brain regions.

**Table 2 (Cont'd)** Fitted Developmental Trajectories of Regional Variability
| Type 2: Nonlinear Increase |  |  |  |  |  |
| --- | --- | --- | --- | --- | --- |
| Caudate_R | 0.72146 | 0.01768 | 2.63E-04 | -475.03 | -475.05 |
| Frontal_Mid_L | 0.70840 | 0.02018 | 1.12E-04 | -477.46 | -477.90 |
| Insula_L | 0.63226 | 0.03143 | 2.84E-07 | -386.63 | -387.32 |
| Precentral_L | 0.71052 | 0.01911 | 4.73E-04 | -425.51 | -425.63 |
| Precentral_L | 0.73864 | 0.01991 | 1.56E-04 | -514.82 | -517.69 |
| Type 3: Linear Decrease |  |  |  |  |  |
| Occipital_Inf_R | 0.83633 | -0.00008 | 4.40E-03 | -445.25 | -443.14 |
| Type 4: Nonlinear Decrease |  |  |  |  |  |
| Temporal_Mid_R | 0.87616 | -0.01227 | 6.18E-03 | -500.10 | -502.79 |
| Temporal_Pole_Sup_R | 0.88247 | -0.01742 | 4.31E-03 | -407.59 | -413.71 |
| Fusiform_R | 0.89809 | -0.01454 | 5.23E-03 | -444.46 | -446.48 |
| Temporal_Mid_R | 0.88328 | -0.01706 | 2.09E-03 | -421.02 | -425.42 |
| Temporal_Mid_R | 0.88400 | -0.01442 | 5.64E-03 | -482.90 | -489.73 |
| Temporal_Inf_R | 0.89785 | -0.01471 | 3.34E-03 | -456.58 | -458.44 |
| Lingual_R | 0.87194 | -0.01768 | 1.10E-03 | -421.13 | -428.85 |
| Temporal_Pole_Mid_L | 0.86929 | -0.01530 | 3.08E-03 | -440.55 | -447.85 |
| Temporal_Mid_L | 0.86986 | -0.01547 | 3.49E-03 | -432.62 | -439.93 |
| Lingual_L | 0.85670 | -0.01517 | 4.56E-03 | -427.97 | -434.71 |

Note: We named each ROI using a traditional naming convention. The same brain ROI names in the table locate at different places representing different sub-divisions of the corresponding (larger) brain regions.

We found that different types of brain regions tend to locate at distinct brain areas. Specifically, the regions with increasing variability mainly located at high-order function-related areas, such as functional association areas in the dorsal and ventral lateral prefrontal areas, the medial prefrontal areas and cingulate cortices, some subcortical regions (putamen, caudate, and insula), and the ventral precentral gyrus. Those with decreasing FC dynamics mainly sit at the visual areas and the temporal lobe. Particularly, for the frontal regions, the ROIs with linearly increased FC dynamics covered a large part of the inferior and superior areas, while the ROIs with nonlinearly increased FC dynamics mainly located at dorsolateral prefrontal areas and ventral precentral areas.

## 4 Discussion

### 4.1 New Findings of the Developmental Patterns of FC Dynamics

To the authors’ knowledge, this paper provides the first-ever early development report of the brain FC dynamics from neonates to two years of age. A recently proposed, robust metric measuring temporal variability of the FC profiles (Zhang et al., 2016; Dong et al., 2018) was adopted to measure such FC dynamics. To comprehensively investigate the development of FC dynamics in the early life, we measured the FC temporal variability at global, functional sub-network, and regional levels. We utilized a longitudinal infant natural sleeping fMRI data with longitudinal data regression model (LMER), instead of cross-sectional data, to better capture the longitudinal developmental trajectories of the FC dynamics. Similar to previous studies on static FC development (Gao et al., 2011; Wen et al., 2019), we also found age-dependent changes in FC dynamics at different levels, including 1) linearly increased global FC dynamics, 2) reduced intra-network FC dynamics in two primary systems and increased FC dynamics in two high-order functional systems, 3) increased inter-network FC dynamics among five functional sub-networks, and 4) a major linearly increasing pattern of the regional FC dynamics in frontal areas.

The first important finding is that the dynamics of FC are reconfigured simultaneously from different scales during the development in the first two years of life. In the large-scale, the whole-brain level FC dynamics is gradually increased (Fig. 2a), indicating an early development towards a *globally* more flexible, adaptive and dynamic functional connectome. Such a pattern was further found to be more complex at a finer, sub-network level. Despite the intra-network FC dynamics did not have any specific developmental patterns for some functional sub-networks, those with decreased intra-network FC dynamics are both primary functional systems (VN and SMN, Fig. 3 and Table 1) and also those with increased intra-network FC dynamics are high-order functional systems (DMN and VAN, Fig. 3 and Table 1). This provides distinctive developmental patterns of FC dynamics among different sub-networks, similar to the previous findings of sub-network-specific static FC development (Gao et al., 2014), where the primary networks are already well developed but the high-order networks are still rapidly developing. For inter-network connectivity, mirroring the previous findings of increasingly static inter-network FC (Gao et al., 2015; Wen et al., 2019), increased inter-network FC dynamics was also revealed (Fig. 3, Table 1). At the regional level, we found more brain regions showing increasing FC dynamics than decreasing ones (Fig. 4, Table 2), despite large heterogeneity in the trajectories of the regional FC variability.

The new findings provide a unique view angle of the rapidly developing brain functions. Higher FC dynamics indicate richer FC patterns, which might indicate more flexibility, higher adaptability, and greater capacity of high-level information processing and integration (Bassett et al., 2011; Zhang et al., 2016; Dong et al., 2018). The developmental changes of high-order function-related networks/areas in FC dynamics thus could explain significantly developed high-level cognitive and behavioral skills in this key period. On the other hand, the biological meaning of the static FC is usually used to support a more static lattice of the brain network, often regarded as a *backbone* or the main strong framework of the brain which is basically less flexible (Biswal et al., 1995; Biswal et al., 2010). Our early developmental findings based on the dynamic FCs could be a good supplement to the previous static FC development studies.

### 4.2 Gradually Increased Global Functional Flexibility in Early Development

Although the FC temporal variability was first calculated for each region, the whole-brain averaging process still revealed a significantly increased pattern from neonates to two years of age. Such a global dominance indicates most of the regions have increased FC dynamics, as both found in a coarse scale (inter-network FC dynamics) (Fig. 3) and a fine scale (Fig. 4). This finding suggests that the infant’s brain is globally evolved to be more and more flexible in early development. Many behavioral studies have presented various emerging high-order cognitive functions in the first a few years of life, such as self-awareness (Amsterdam, 1972), spatial attention (Haith et al., 1988), and working memory (Reznick, 2008). All these skills request better cognitive flexibility, controllability, adaptivity for brain regions to accommodate and adapt to quickly changing environments (Medaglia et al., 2018). For example, to respond to the internal (i.e., feelings about self) and external stimuli (i.e., multimodal sensory input from the environment) promptly, the task control and attention control regions in the infant brains are more likely to become increasingly variable with increased FC flexibility, i.e., their FC profiles could be more variable, linking different functional systems at different time for their coordination and communication.

Our findings are also informative when comparing with the previous studies on different age groups (mostly older age groups) trying to link brain functional flexibility to the behavioral performance or outcome. Mounting evidence has been indicating that human brain dynamics are important for establishing cognitive abilities and behaviors during childhood and adolescence development (Bassett et al., 2011; Bassett and Mattar, 2017). For example, Bassett et al. found that the brain flexibility is positively related to the human learning ability in order to adapt existing neural processes and brain functions to derive new patterns of the desired behavior (Bassett et al., 2011; Bassett and Mattar, 2017). Medaglia et al. (2018) identified the increasing dynamics of “brain states” that were associated with improved executive function during the development at early adulthood (8-22 years of age). It is thus highly needed to have more future studies to investigate the relationships between brain dynamics’ changes and behavior development in the first two years of life.

The more flexible global FC along with aging found in our study also provides evidence for previous long-standing hypothetical models, i.e., there exists continuously increased brain functional organization *efficiency* at the global level. In Cao et al. (2017), from a connectome (topology of the entire brain static FC) perspective, they proposed that the brain networks are changing towards an “organized and optimized” configuration, with a “strengthening global *balance* between local and global information processing”. That is, the local (averaged across the whole brain) and global efficiency (reflecting functional segregation and integration, respectively) are increasing at an equal rate. Similar hypothetic models of local and global efficiency development are also reported in another review paper of early FC development (Zhang et al., 2018) and another study based on a longitudinal infant fMRI data from a modular perspective (Wen et al., 2019). Another early fMRI study directly investigating local and global efficiency development in the first two years after birth, however, shows a significant increase in both metrics from neonates to one year old, but such increasing pattern seems stopped (and even slightly reduced for local efficiency) from one to two years old (Gao et al., 2011). Studies on the whole-brain structural connectome revealed increasing integration but decreasing functional segregation, possibly due to different definition between FC and structural connectivity (Yap et al., 2011). Nevertheless, all these previous studies on the efficiency development are based on region-wise paired relationship and topological analysis of *static* FC, mostly *stronger* ones, which generally measures the “backbone” of the connectome. To better characterize “efficiency”, we propose a *temporal dynamics perspective* of efficiency, which further defines regional flexibility (the FC profile between each region to all other regions changes along time). Such temporal flexibility may create a transient “short-cut” from one functional system to another, despite their weak static FC, to facilitate information flow and exchanges. From the dynamic perspective, we again validate the hypothesis that there is a globally increasing trajectory of the connectivity efficiency (in terms of flexibility) but we also reported a new finding that such a flexibility change has a linearly increasing pattern, indicating that the FC flexibility is still under a fast increasing track and does not reach its peak at two years of age.

### 4.3 Different Developmental Trajectories of Intra-network FC Flexibility between Primary and High-order Sub-networks

Our findings at the sub-network level (decreased *V*_within-net_, for primary functional sub-networks but increased *V*_within-net_ for high-order functional sub-networks) further validate the previous findings of different developmental trajectories between primary (especially for primary visual and sensorimotor functions) and high-order functional systems. Specifically, based on the static FC, previous findings consensus on much earlier maturing rate for these primary functional networks than the high-level ones (Gao et al., 2014; Gao et al., 2015), which explains the order of the behavioral development. Our study provides a new evidence of such developmental differences from a dynamic perspective, providing some new knowledge about the development at this intermediate level.

Rather than measuring the properties of the static FCs in each sub-network, *V*_within-net_ evaluates how within-network FC fluctuates across time, where a larger *V*_within-net_ value indicates richer FC patterns possibly underlying more frequent information communications among *different components* in the same functional systems (Dong et al., 2018; Sun et al., 2018). We have known that certain high-order functional systems in the adult population often constitute sub-systems in order to mediate more complex functions. For example, the DMN was found to have multiple functional sub-networks, each of which may be responsible for distinctive functions (Buckner et al., 2008; Assaf et al., 2010; Li et al., 2013). A meta-analysis study with task-based activities found that the DMN is functionally heterogeneous, corresponding to different behavioral metadata and indicating that the DMN is differentially specialized (Laird et al., 2009). With temporal lag-respected task fMRI source separation, four different DMN sub-networks were revealed, each with different spatiotemporal relationship to different cognitive components (Van De Ville et al., 2012). A recent dynamic FC study conducted a clustering analysis on transient co-activations of rs-fMRI signals and revealed multiple DMN-related co-activation patterns, each of which indicated a transient connection between the DMN and other regions from other networks (Liu and Duyn, 2013). Such spatiotemporal flexibility of the DMN was even observed in a sustained attention task (Li et al., 2015). Collectively, the coordination and interactions among different DMN sub-networks are important for the infant to accomplish more complex tasks, which could be reflected by the increasing *V*_within-net_ of the DMN. In the same vein, the VAN (or equivalently, salience network) has also been suggested to have different sub-systems (Chand et al., 2017) and could be heavily involved in mediating the switch between DMN and FPN (Sridharan et al., 2008; Menon, 2011; Nekovarova et al., 2014). Thus, we speculated that the intra-network FC within DMN and VAN is becoming more flexible as their respective sub-system structures become much clearer along the development to facilitate high-level cognitive functions.

The reversed developmental trajectories of *V*_within-net_ of the VN and SMN (the only two primary functional sub-networks) may, on the other hand, indicate increasingly homogeneous FC architecture of each network along the development. It is reasonable because the previous studies have found that the brain functional sub-networks are generally defined according to the anatomical closeness at birth, but with long-term FC becoming stronger, they are gradually developed to be spatially distributed (Yap et al., 2011; Cao et al., 2017; Wig, 2017; Gilmore et al., 2018). As the VN and SMN atlases are defined according to an adult-based parcellation, they may not well suit for neonates (also include some regions from other functional sub-networks), leading to greater variability in their respective within-network FC. With the increase of age, the VN and SMN become more and more similar to the adult pattern with more focal spatial distribution and thus generate a smaller FC variability, reflecting the highly specialized function of each sub-network. Another possibility is that the increasing myelination within the primary networks makes the intra-network FC within the VN and SMN much stronger, leading to the less freedom for these FCs to be fluctuating. Such stability of their FC could be due to the strong within-network FC backbone, which have the ceiling effect, making the FC fluctuation less possible (Bassett et al., 2013). On the other hand, the decreasing flexibility could indicate that the primary sensory functions require more stable FC to maintain a stable and robust bottom-up information feeding to high-level functional sub-networks. Yet, such decreasing *V*_within-net_ of the VN and SMN seems to follow nonlinear (exponential) trajectories, meaning their *V*_within-net_ turn to be smaller but such changes are becoming less and less until the *V*_within-net_ are stable in later ages. Such earlier stabilized *V*_within-net_ changes in the primary function-related sub-networks vs. the possibility of prolonged changes in the high-order functional systems again shows fundamental differences between two types of functional systems.

### 4.4 Increased Inter-network FC Flexibility

We found that nearly all the significantly changed *V*_between-net_ are linearly increasing (more flexible and diverse) with age. If not counting the LN/SN and VN, all the *V*_between-net_ among the rest of five sub-networks (SMN, DAN, DMN, FPN, and VAN) are increasing with age. This indicates that inter-network information communication becomes more frequent and more active in the first two years of age. For example, the VAN, FPN, and DMN that are involved in a “triple network” hypothetic model (Menon, 2011) showed linear increases in their pairwise *V*_between-net_. It has been proposed that such three networks are the “core” neurocognitive networks and their functional interactions are important for complex, high-order cognitive functions (Menon, 2011) and could be responsible for psychiatric and neurological disorders, such as depression (Berman et al., 2011), schizophrenia (Palaniyappan et al., 2011), and dementia (Zhou et al., 2010; Yu et al., 2017). Recently, researchers have found that the dynamic network switch supports flexible attention (mainly via DAN and VAN) and cognitive control (mainly via FPN) to meet time-varying changes in cognitive demands, and the loss of such optimized dynamics may lead to poor task performance in a decision-making task (Taghia et al., 2018). Such a flexibility among the three networks could be associated with Attention-deficit/hyperactivity disorder (ADHD) (Cai et al., 2018) and schizophrenia (Supekar et al., 2019). Similarly, another study from the “modular” perspective also found the relationships between the three core networks and the learning ability (Bassett et al., 2011) in healthy subjects and with the pathophysiology of schizophrenia (Braun et al., 2016). It is worth noting that a previous static FC-based study has identified significant changes in the averaged inter-network FC strength between the VAN and DMN and between the FPN and DMN, but not between the VAN and DMN (Gao et al., 2015). Our finding of all increasing *V*_between-net_ among the three core networks gives the first evidence that interactions among these three networks are also very important for supporting the remarkable development of complex cognitive abilities over the course of development during early infancy. The use of dynamic FC further increases the sensitivity in the detection of developing inter-network functional interactions. As such, all high-order function-related sub-networks have increased *V*_between-net_ in the first two years of age.

Another interesting finding is that the SMN have increasing *V*_between-net_ with all the other four high-order cognitive function-related sub-networks (DAN, DMN, FPN, and VAN) but had decreasing *V*_within-net_ itself. We tentatively interpreted such a result as differently developed functional integration and segregation in a dynamic viewpoint. For within-SMN FC, the connectivity becomes stable while gaining its strength to make sensory inputs and motor outputs interact and coordinate better (more stable) and less affected by other factors. However, the input sensory information needs to be spread and further processed in high-order functional sub-networks such as DAN, DMN, FPN, and VAN, and the motor performance could be highly modulated and adaptively controlled by these high-order functional sub-networks (Hutchison et al., 2013; Calhoun et al., 2014). During early development, the increasingly adaptive ability of the brain is to better fit for or response to the moment-to-moment changes in the environment, as well as time-varying attention and task control demands during the conduct of complex cognitive functions, all of which requires a timely response and continuous preparedness at different attention levels. The increasing *V*_between-net_ of the SMN suggests richer FC patterns underlying feed-back and feed-forward interactions between it and other high-order functional sub-networks (Dong et al., 2018), which may allow infants to carry out more and more complex tasks (Reddy et al., 2018). Combing with the heightened stability (decreased *V*_within-net_) of FC within the SMN, we speculated that the increased dynamic reconfigurations between SMN and high-order networks may due to the enhanced ability of infants for integrating internal and external sensory and perception input to achieve the rapid development of complex functions.

### 4.5 Heterogeneous Developmental Trajectories of Brain Regions’ FC Dynamics

The measurement of the nodal FC variability at the finest scale provides us a more detailed picture of the rapidly developing brain dynamics. As shown in Fig. 2b (cross-sectional) and Fig. 4 (longitudinal), convergent evidence shows complex spatiotemporal changes in the flexibility of the FC architecture across ages. We found heterogeneous developmental trajectories (types 1–4) for different regions, where those with linearly increasing FC flexibility were dominant to all sub-networks but VN (to which nonlinear decrease is dominant) and FPN (to which significant more nonlinear increased type-2 regions were found). Generally, those with decreasing FC flexibility located at the inferior part of the brain, while those with increasing FC flexibility were widely distributed at the anterior and superior part of the brain. This is consistent with the previous consensus of the developmental order of the brain function, that is, from posterior to anterior, from inferior to superior, from medial to the lateral areas (Gao et al., 2014).

Specifically, the frontal and parietal association areas and the projections from them to the subcortical regions could have protracted maturation process (Zhang et al., 2018). Many of these regions were found to be developed from non-hubs (the regions with fewer connections) or provincial hubs (the regions with more intra-network connections) to connector hubs (the regions that simultaneously connecting two or more functional sub-networks) (Wen et al., 2019). These regions may serve as a bridge for information exchange and integration among different systems. With development, their roles could become more and more important. This is also consistent with the consensus of developmental neuroscience, that the frontal areas mediate more complex functions and develop last. Type-3 developmental trajectories only occur at the temporal fusiform area that is for face perception (Kanwisher et al., 1997) as an important node in a ventral “what” visual pathway (Hickok and Poeppel, 2007). In our previous work on developmental modular changes, we also found that this region might develop from non-hub to provincial hubs, making its FC more stable (i.e., less flexible).

Additionally, we found that the number of brain regions with linearly increased FC variability (50 ROIs) is much more than those with nonlinearly increased variability (8 ROIs), suggesting that most of the complex cognitive functions have protracted development at later ages. The frontal areas are generally split into two categories, the type-1 category (linearly increase) mainly covered the inferior and the superior frontal areas (with functions like self-awareness (Goldberg et al., 2006) and motor planning (Grier, 2005)), while the type-2 category (nonlinearly increase with earlier saturation) mainly sit at the ventral precentral areas (mouth, tongue, and laryngeal movements). These results suggest that the language-related motor functions may have been largely in place by the end of two years old, while other more complex functions such as language production, movement preparation, initialization, and control could continue to be developing at later ages.

## 5 Conclusions

In this study, we provided a first-ever comprehensive report of how brain functional dynamics develop in the first two years after birth at the different spatial scales. Our results indicate that the brain is becoming increasingly flexible, dynamic and adaptive with complex spatiotemporal developmental patterns. The unraveled detailed changes in the brain functional dynamics pave a road for the future works for a better understanding of infant brain development in the chronnectome perspective.

## Acknowledgments

This study was supported in part by National Institutes of Health (MH100217, MH108914, MH107815, EB022880, AG041721, MH110274, MH116225, and MH117943). This work utilizes approaches developed by an NIH grant (1U01MH110274) and the efforts of the UNC/UMN Baby Connectome Project (BCP) Consortium. This work also uses the data from “Multi-visit Advanced Pediatric brain imaging study for characterizing structural and functional development (MAP Study)”.

